# A haplotype-aware de novo assembly of related individuals using pedigree graph

**DOI:** 10.1101/580159

**Authors:** Shilpa Garg, John Aach, Heng Li, Richard Durbin, George Church

## Abstract

**Motivation:** Reconstructing high-quality haplotype-resolved assemblies for related individuals of various species has important applications in understanding Mendelian diseases along with evolutionary and comparative genomics. Through major genomics sequencing efforts such as the Personal Genome Project, the Vertebrate Genome Project (VGP), the Earth Biogenome Project (EBP) and the Genome in a Bottle project (GIAB), a variety of sequencing datasets from mother-father-child trios of various diploid species are becoming available.

Current trio assembly approaches are not designed to incorporate long-read sequencing data from parents in a trio, and therefore require relatively high coverages of costly long-read data to produce high-quality assemblies. Thus, building a trio-aware assembler capable of producing accurate and chromosomal-scale diploid genomes in a pedigree, while being cost-effective in terms of sequencing costs, is a pressing need of the genomics community.

**Results:** We present a novel pedigree-graph-based approach to diploid assembly using accurate Illumina data and long-read Pacific Biosciences (PacBio) data from all related individuals, thereby generalizing our previous work on single individuals. We demonstrate the effectiveness of our pedigree approach on a simulated trio of pseudo-diploid yeast genomes with different heterozygosity rates, and real data from *Arabidopsis Thaliana*. We show that we require as little as 30× coverage Illumina data and 15× PacBio data from each individual in a trio to generate chromosomal-scale phased assemblies. Additionally, we show that we can detect and phase variants from generated phased assemblies.

**Availability:** https://github.com/shilpagarg/WHdenovo

## 1 Introduction

The ability to faithfully reconstruct genomes is a crucial step in better understanding evolution and the nature of inherited disease (Tewhey *et al.* (2011)). *De novo* genome assembly aims to address this problem by generating complete genome sequences from error-prone sequencing reads alone, without the use of a reference genome. Creating a *de novo* assembler that resolves genomic repeats and is generalized to genomes of varying heterozygosity rate (the average proportion of loci that differ between homologous sequences) has posed a significant challenge to the scientific community. Assembling diploid genomes adds further difficulty; in order to accurately represent diploid genomes, assemblies must correctly identify and *phase* (i.e. determine the correct haplotype of) homologous sequences. One promising approach to diploid genome assembly is incorporating sequencing information from a related set of individuals, particularly from mother-father-child trios, and using the Mendelian information offered by the corresponding pedigree to infer the layout of alleles along homologous sequences.

Some assemblers (Levy *et al.*, 2007; Pryszcz and Gabaldón, 2016; Simpson and Durbin, 2012; Bankevich *et al.*, 2012; Li, 2015) tackle single-individual assembly using Next-Generation Sequencing data; however, while accurate, the short length of NGS reads often leads assemblies to fragment at repetitive and highly-heterozygous regions. Other assemblers (Koren *et al.*, 2017; Vaser *et al.*, 2017; Berlin *et al.*, 2015; Chin *et al.*, 2013) utilize longer Third-generation sequencing reads to obtain more contiguous sequences, and to help resolve repeats and heterozygous regions. Yet, these require high coverage due to the high error rate in long-read data, which is very costly. *Hybrid* assemblers utilize both types of reads, taking advantage of the accuracy of short reads and the scale of long reads to generate complete, high-quality assemblies (Bashir *et al.*, 2012; Antipov *et al.*, 2015; Zimin *et al.*, 2017).

However, the methods described above collapse differences between homologous pairs into a single consensus sequence, without regards for the rich information given by the layout of different alleles along two DNA strands (Simpson and Pop, 2015). By contrast, other assemblers by Chin *et al.* (2016); Garg *et al.* (2018); Weisenfeld *et al.* (2017) have been developed to generate *haplotigs*, haplotype-resolves assemblies for diploid genomes.

Certain assemblers have been developed to employ pedigree information into the process of assembly as well, specifically for the case of mother-father-child trios. For example, trio-sga creates haplotypes of the child based on parental Illumina data (Malinsky *et al.*, 2016), while TrioCanu uses such data to partition child long-read data and subsequently assembles the partitioned reads separately (Koren *et al.*, 2018). Yet, these two methods cannot phase variants which are heterozygous in all three individuals in a trio, and by relying solely on parental Illumina data, may not correctly haplotype long reads which cover repetitive genomic regions. Furthermore, TrioCanu does not work properly at low coverages of long-read data.

Another method approaches the diploid assembly problem by aligning long-read data from all individuals in a pedigree to a reference-genome, then finding the most likely partitioning of reads as determined by the PedMEC problem (Garg *et al.*, 2016). Essentially, the PedMEC (Weighted Minimum Error Correction on Pedigrees) problem finds the partitioning of reads from related individuals that incurs the least cost, which is calculated based on the likelihood of errors occurring at various locations along the reads as well as recombination costs between each site of heterozygosity. This method, however, only concerns bi-allelic variants, and contains reference bias, meaning that unique DNA sequences may not be detected or phased correctly.

Thus, developing a haplotype-resolved de novo assembly approach for related individuals which is cost-effective, flexible with regard to genomic complexity and heterozygosity rate, and which does not contain reference bias, is a pressing need for the genomics community.

### Contributions

Our graph-based method, implemented as a new tool WHdenovo, performs phasing in the space of a *pedigree graph* (defined below), and is generalized to assemble genomes of varying heterozygosity rate and with multi-allelic variants, thus allowing for the creation of accurate, complete haplotigs.

More precisely, our approach builds a pedigree graph, or joint sequence graph, using combined Illumina data from all individuals in a pedigree; the graph represents heterozygous locations as *bubbles*, as shown in Figure 1. Given a pedigree graph containing a series of bubbles, long-read (PacBio) data from each individual are threaded through it; in essence, the most probable paths that the long reads trace through the bubbles in our pedigree graph, and which obey the Mendelian constraints imposed by the pedigree, represent our true haplotypes. We draw key concepts from the graph-based assembly approach for single individuals described in Garg *et al.* (2018) and the PedMEC formulation set forth in Garg *et al.* (2016), yet synthesize and extend them to overcome their respective shortcomings.

**Fig. 1.**
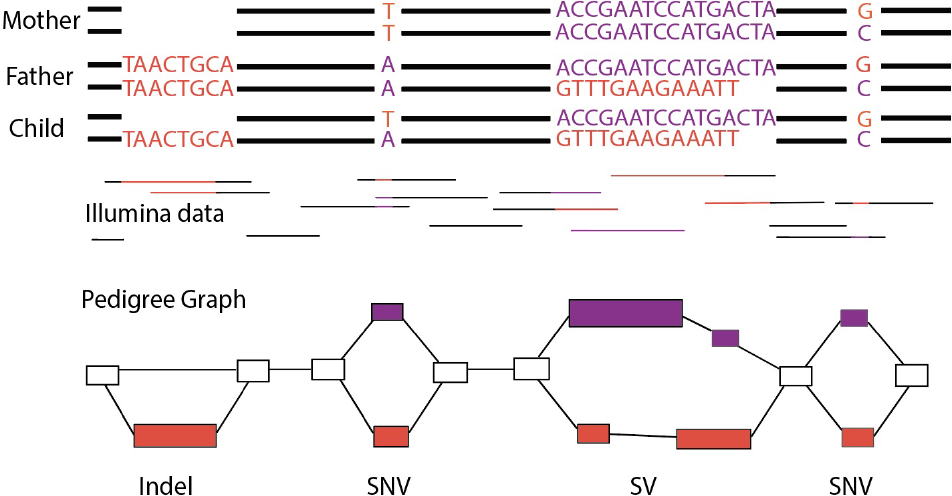
The Illumina data (middle) from the trio of genomes can be represented as a pedigree graph. The bubbles in the graph (bottom) show four different variants; from the left, there is an indel, SNV, SV, and SNV.

Our graph-based approach poses several advantages. For example, we can detect and phase all types of small and large variants, and require relatively low coverage of costly long-read data. We can also phase variants that are heterozygous in all individuals; for example, SNV1 from Figure 2. Moreover, by incorporating hybrid data from all related individuals, we can effectively phase reads in repetitive genomic regions, and as shown by SV2 in Figure 2, if parental reads span a variant but child reads do not, we can still correctly identify and phase the variant in all three individuals. Assemblers such as TrioCanu would not be able to phase variants under these circumstances, and may forced to break assembly contiguity.

**Fig. 2.**
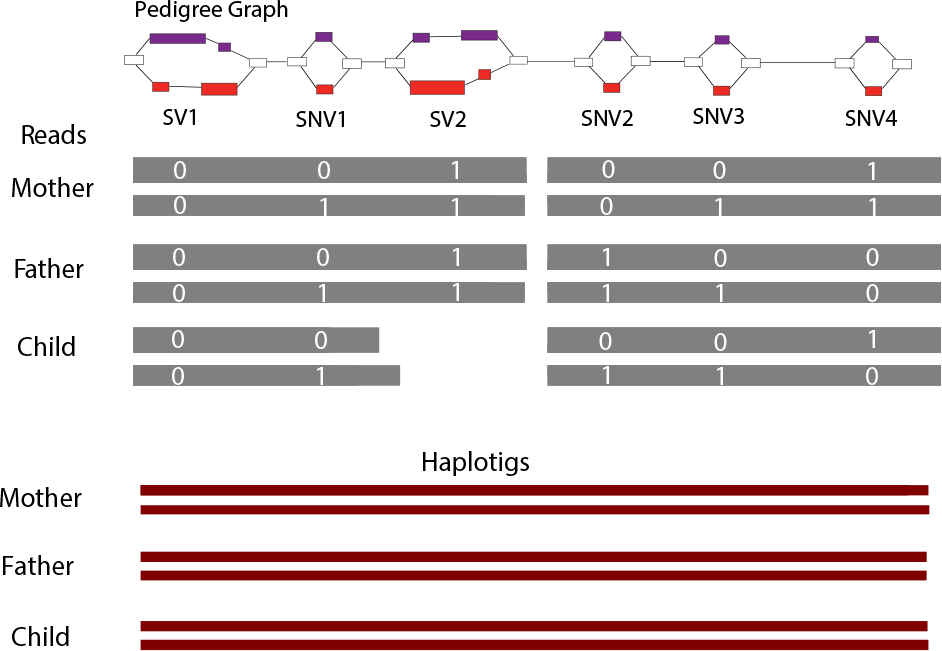
Input: This figure shows the pedigree graph (top) (consisting of four SNVs and two SVs) and the PacBio reads (gray) with respective alleles (white digits). Output: the final haplotigs (crimson) for each individual in a trio. Our method can phase the variants that are heterozygous (SNV1) in all individuals and any variant covered by atleast one read from any individual (SV2 in child), resulting in continuous and complete haplotigs

To demonstrate the practical effectiveness of our method of haplotype-aware *de novo* assembly for related individuals, we assemble two sets of genomes. First, we haplotype the genomes of a simulated trio of pseudo-diploid yeast, which allows us to comprehensively study assembly at varying read coverage and heterozygosity rates. Then, we use real data to assemble the complete diploid genome of a trio of *Arabidopsis Thaliana*. These results indicate that our hybrid method is adaptable to genomes of varying heterozygosity rates. We demonstrate that our method is cost-effective, requiring only 30× short-read coverage and 15× long-read coverage for every individual in a pedigree to generate near chromosomal-scale assemblies for all individuals. Moreover, at these coverages, we show that our assemblies for both real and simulated data are more accurate and contiguous when compared to those produced by TrioCanu at 45× child long-read coverage. In a final experiment, we also show that we can detect and phase variants.

## 2 Pedigree-aware phased assembly pipeline

In this section, we present the workflow of our pipeline, which takes input as raw Illumina and PacBio sequencing data from all related individuals in a pedigree, and outputs final, polished haplotigs. Once we create a pedigree sequence graph, our goal is to find the walks through this graph that correspond to the true haplotypes of all related individuals. These *haplotype paths* will encode the phasing of all variants in the haplotigs for each individual in the pedigree. Due to errors of Illumina data and genomic characteristics such as repeats, there are inevitably multiple paths through this graph that do not correspond to true haplotype paths. Thus, to construct the true haplotype paths, we seek the maximally likely paths based on confidence scores of how the nodes are connected to each other over long distances, which we determine using PacBio reads aligned to our graph.

This pipeline generalizes our previous single-individual approach (Garg *et al.*, 2018) to related individuals to yield the chromosome-scale haplotigs. Figure 3 represents a conceptual workflow of our pipeline, detailed below.

**Fig. 3.**
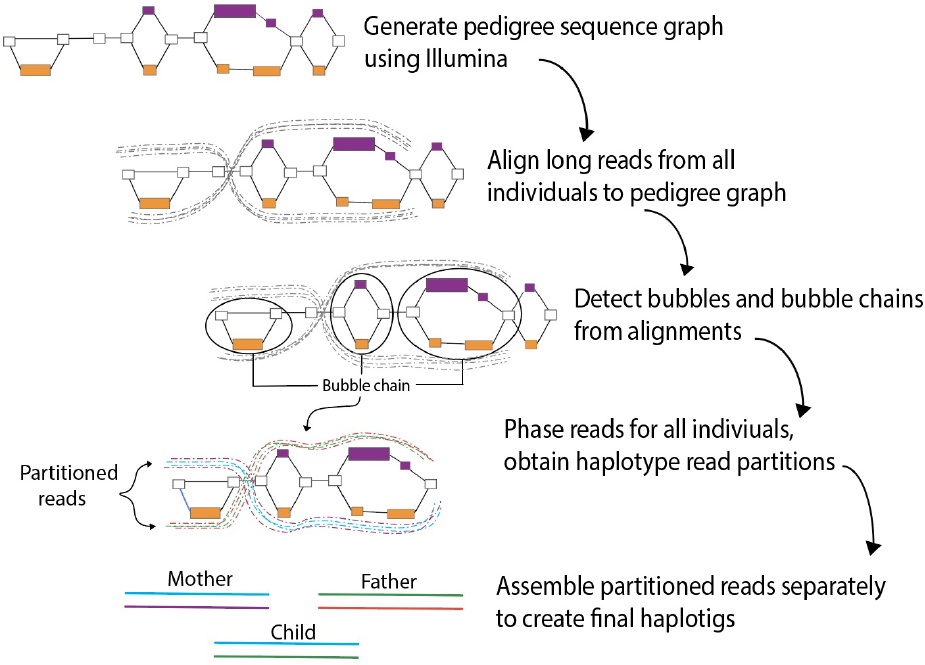
Overview of the pedigree-aware phased assembly pipeline. After we generate a pedigree graph using Illumina data, we align PacBio reads from all related individuals from a pedigree to the graph. Using these alignments, we detect ordered sequences of heterozygous regions, represented as bubble chains. We then find the best partitioning of each set of reads into haplotypes based on the paths taken through the bubble chains, and assemble the partitioned reads.

### Pedigree Graph

We use short, accurate Illumina reads from all related individuals as the basis for generating a pedigree graph. We denote the bidirected pedigree graph as *G*_*p*_, containing a set of nodes *N*_*p*_ and a set of edges *E*_*p*_. Conceptually, each node *n*_*i*_ ∈ *N*_*p*_ represents a segment of DNA found in the Illumina data, and the node 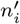 represents its reverse complement. Note that because nodes are generated using the combined reads of all individuals in a pedigree, not every node sequence is necessarily present in the genome of every individual. Additionally, nodes can be traversed in either direction; when traversed in the reverse direction, its sequence is considered reverse-complemented. Every edge *e*_*ij*_ ∈ *E*_*p*_ represents an adjacency between the sequences represented by node *n*_*i*_ and *n*_*j*_.

The pedigree graph *G*_*p*_ contains a set of bubbles *L* that represents heterozygous variants (or sequencing errors). The terminology *bubble* is drawn from Garg *et al.* (2018) and follows from the *ultrabubble* coined by (Paten *et al.*, 2017). Graphically speaking, bubbles are directed and acyclic, biconnected, and minimal, as discussed by Garg *et al.* (2018). We define each bubble *l*_*k*_ to be the set of allele paths contained within it, where each allele path is a unique sequence of nodes spanning a common start and end node. Intuitively, each bubble is bookended by common sequences —the start and end nodes —and the unique node sequences connecting them represent the alleles gleaned from our Illumina data. Figures 1 demonstrates the bubbles representing various variants.

### PacBio Alignments

We align PacBio long reads from all individuals to produce paths through the pedigree graph *G*_*p*_, in a similar process to Garg *et al.* (2018). The concept of *aligning* a PacBio read to *G*_*p*_ captures the idea that the sequence of a single PacBio read can trace a path through the sequences contained in many nodes. Thus, for a given PacBio read, we define a read alignment *r*_*i*_ to be a path through *G*_*p*_, defined by the oriented nodes *n*_1_ … *n*_*k*_ which map to a particular read. Given a set of individuals 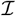, we will align the PacBio reads from every individual 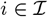 to *G*_*p*_, resulting in a set of read alignments *R*_*i*_ = {*r*_1_, *r*_2_, …, *r*_*j*_} for every individual. As an intuitive checkpoint, note that as PacBio reads from different individuals trace different paths through nodes and bubbles in *G*_*p*_, we gain information about not only the correct ordering of nodes to form the genome in question, but also the alleles and phasing present in each individuals.

### Bubble Chains

To consolidate the information gained by the PacBio alignments, we need to formalize the process of finding ordered *bubble chains* which represent the layout of heterozygous sites across the genome. Ideally, we would be able to use our PacBio alignments to obtain *C*, a set of bubble chains containing one continuous bubble chain corresponding to each chromosome. Before generating bubble chains, we must perform a series of filtration steps to remove erroneous bubbles according to the algorithm described in the following paragraph:

On input 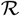, the set of all read alignments *R*_*i*_ for individuals 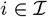, we project all partial alignments to bubble space—that is, for every instance where a read traverses a bubble, we replace the corresponding set of nodes by the appropriate bubble ID. We now perform our filtration steps; for every pair of nodes or bubbles from every alignment path in 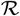, denoted as 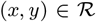, we compute the coverage *c*_(*x,y*)_, and if *c*_(*x,y*)_ < 5, we remove the pair. Next, we calculate the degree of every remaining node *x*, and if *deg*(*x*) ≥ 3, we remove all pairs containing *x*. Using the resulting filtered pairs, we then perform DFS to find *U*, an initial set of unambiguous bubble chains termed *unitigs*. Finally, if there is at least one read connecting two unitigs every pair of unitigs in *U*, we can gain information about the ordering between these initial unitigs. Record the resulting orderings as *C*, the set of final bubble chains. Figure 4 demonstrates the algorithm with a basic example.

**Fig. 4.**
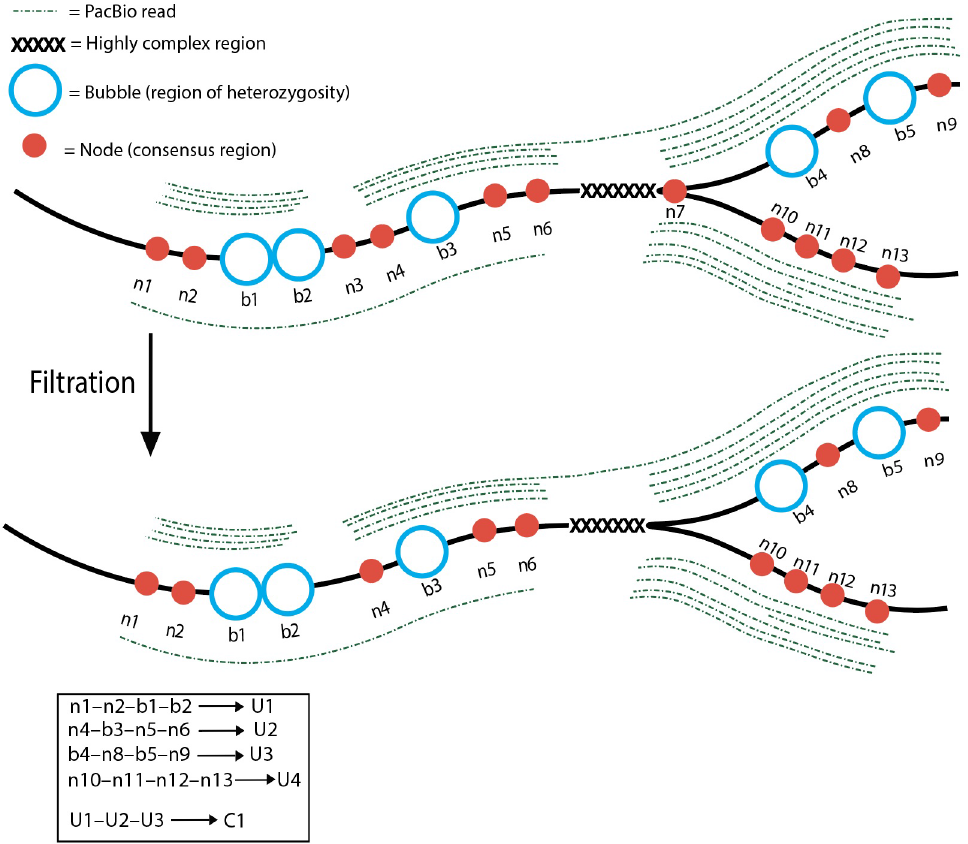
This figure considers a toy example of our algorithm to find bubble chains. Shown is the joint sequence graph, containing nodes and bubbles, with many reads spanning them. In the filtration step, n3 is filtered because it is only covered by a single read, and n7 is removed because it branches (has degree > 2). Unitigs U1-4 are generated by using DFS for the pairs of edges and bubbles whose connection are covered by at least 5 reads. Because there is a read that spans U1 and U2, and a read that spans U2 and U3, we can combine these smaller unitigs into larger final bubble chain C1.

### Graph-based phasing on pedigrees

We introduce the *Graph version of Weighted Minimum Error Correction on Pedigrees Problem*, or gPedMEC, as our central phasing algorithm. gPedMEC relates to the PedMEC formulation set forth by Garg *et al.* (2016), which concerns alignment to a reference genome and applies only to bi-allelic variants. Considering multi-allelic variants in the gPedMEC framework requires additional phasing-related observations. Specifically, bubbles which represent biallelic variants ensure that a child’s haplotype paths must be the same as a combination of parental haplotype paths. By contrast, in bubbles containing multi-allelic variants, it is possible for child haplotype paths to differ from parental ones; this would occur under the circumstance that the variant encoded by the bubble is long, and a new allele were created as a result of recombination within the variant itself. Based on these insights, we create a new algorithm to determine the haplotypes of any set related individuals in a more flexible, accurate, and representative manner.

Ultimately, the goal of solving gPedMEC is to recreate the haplotype paths of every individuals through the bubble chains we calculated in *C*. The haplotype paths for a given individual are determined by deducing the two paths through the bubble chains which incur the least cost (i.e. are most likely), where cost is determined by confidence in alignment paths and recombination costs. Doing so will inherently also compute long-read partitionings.

In order to represent the paths taken through our bubble chains for each set of alignments *R*_*i*_, we create *bubble matrices* 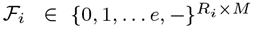 for each individual *i*. Here, *e* is the maximum number of alleles contained in any bubble, and *M* is the number of bubbles in a chain. Figure 5 shows a sample 3-bubble chain with PacBio reads from each individual aligned to it; the corresponding bubble matrices are shown for the three individuals are shown below:

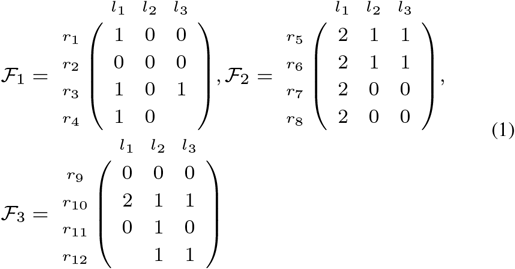

**Fig. 5.**
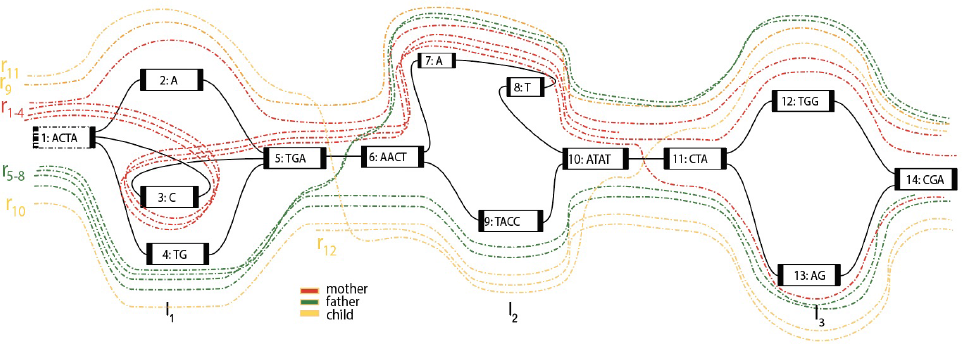
For a subgraph of *G*_*p*_, the example shows three bubbles *l*_1_, *l*_2_, and *l*_3_, and their corresponding alleles. Reads from mother, father and child traverse the bubbles.

If all reads in a bubble matrix 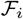 were to follow exactly one of two paths through a bubble chain, solving for the true haplotype paths for the corresponding individual, 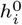, 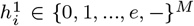, would be trivial. For example, 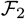 is conflict–free, and the corresponding haplotype paths are self-evident. However, due to long-read sequencing errors, conflicts within bubble matrices are inevitable. Thus, our goal is to find the optimal set of matrix entries which can be flipped in order to create conflict free matrices containing a bipartition of reads which follow the two haplotype paths of each individual (and which obey the Mendelian constraints of the pedigree). So, for every individual, we also need to introduce a weight matrix 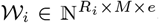, where each entry 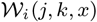 represents the cost of flipping read *j*, at bubble *l*_*k*_, to allele *x*, where *k* ∈ {0, 1, …*M*} and *x* ∈ {0, 1…, |*l*_*k*_|}. An example weight matrix, corresponding to 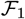, is shown below:

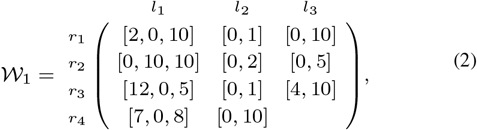

Other weight matrices can be written similarly.

We need to account for the Mendelian constraints imposed by the pedigree as well. Thus, we introduce the *transmission vectors* 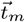, 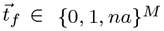 for each triple 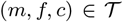 to denote the alleles passed from each parent to child at every location in a bubble chain, where 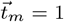 if the mother passes on an allele from 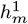, and so on. Under the circumstance that recombination occurs *within* a bubble, in which case the value of *na* is passed on, no haplotype of any parent is directly transmitted to the child. Thus, the recombined sequences will form allele-paths in the bubble.

Additionally, we introduce recombination costs 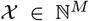, where 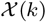 denotes the cost between bubbles of indices *k*–1 and *k*. For example, if a transmission vector 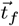 passes on an allele from 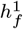 at location *k* – 1 and an allele from 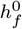 at *k*, this would not incur any recombination cost 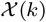. Yet, when transmission vector 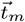 passes on *na* at location *k* – 1 and an allele from 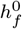 at bubble *k*, this would not incur any recombination cost.

Having defined all necessary terms, we can now more clearly define the gPedMEC problem. Consider a pedigree graph *G*_*p*_ containing bubble chains *C* and recombination costs 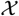, from a pedigree with individuals 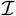 and relationships 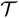; each individual has corresponding PacBio read alignments *R*_*i*_, and matrices 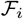 and 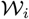. The crux of gPedMEC is then to determine the set of all bubble matrix entries to be flipped which accrues the minimal cost, based on a) the weight of flipping these entries coupled with b) the recombination cost incurred by the transmission vectors implied by the resulting child haplotype paths. Once these matrix entries are determined, the haplotypes 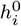 and 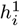 for all individuals become self-evident.

#### Dynamic Programming to Solve gPedMEC

In the following section, we discuss our dynamic programming (DP) algorithm to solve gPedMEC, which is the generalization of Garg *et al.* (2016), to which we refer the reader to for additional information about the algorithm. The motivation behind using a dynamic programming algorithm is utilize a DP table to determine the optimal haplotype paths more efficiently than with a brute force algorithm, which would require exponential time with respect to the number of bubbles and alignments.

#### DP cell initialization

As in Garg *et al.* (2016), we proceed by first computing the initial DP cell cost Δ_*C*_ (*k, B, t*) accrued by flipping entries in column *k* of each bubble matrix, where *k* is the index of bubble *l*_*k*_, *B* is a bipartition of reads and *t* ∈ {0, 1, *na*}^2^ is a transmission tuple indicating which haplotype from mother and father is respectively inherited, if applicable.

Intuitively, there is a relationship between a bipartition *B*, a transmission tuple *t*, and the resulting set of readsets induced at location *k*, termed 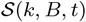. For example, if the transmission tuple does not contain any *na* values, then the child’s haplotype path *must* be a combination of parental haplotypes - correspondingly, the reads in the child’s bubble matrix can be merged with bipartitions in the parental bubble matrices due to the identity by descent (IBD) condition. In the case of a trio, this would lead to two resulting bipartition readsets. In the presence of *na* values, the child’s reads would not be merged with parental reads, leading to three bipartition readsets in the case of a trio.

For a given bubble *l*_*k*_, each bipartition can be assigned any pair of alleles (*x, y*), of which there are 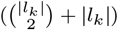 (the added |*l*_*k*_| accounts for the possibility of assigning the same allele). To notate this, we let 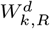 represent the cost of flipping reads for a specific bipartition readset 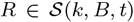 to an allele-pair (*d*_1_, *d*_2_) ∈ *A*, where A is the set of allele-pairs {(*x, y*) ∈ *l*_*k*_ × *l*_*k*_|*x* ≤ *y*}:

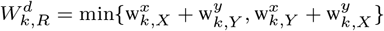

where *X* and *Y* are the readsets of a bipartition readset *R*. The cost of flipping to a given allele can be computed as:

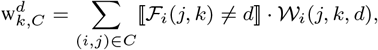

Thus, to initialize the cost Δ_*C*_(*k*, *B*, *t*), we need to find the assignment of allele pairs to all bipartition readsets which incurs the minimal flipping
cost. This can be formalized in the following way:

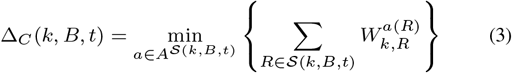

The inner sum computes the minimum allele-pair assignment cost from all bipartitions in 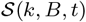. The calculations in Figure 6 provide a more concrete example for calculating DP cell initialization cost. In this example, we assume a sample partition of reads and child partition as per transmission value shown in Figure 6(c) where the mother passes on the green allele and the father passes on the blue. To calculate the DP cell initialization cost in (d), we find the assigment of allele-pairs to each of the bipartition readsets which incurs the lowest cost. For example, the cost associated with the allele-pair assignment (0,2) and (0,1) to bipartitions (green, purple) and (orange, blue), respectively, is (2) + (5+8) + (4+1) + (3+4+4) = 29 because the minimum cost of flipping all reads in the green partition to allele 0 is 2, that of flipping the purple partition to allele 2 is (5+8) = 13, and so on. In this way, we can compute costs for other allele-pair assignments and store the minimum in the DP cell.

**Fig. 6.**
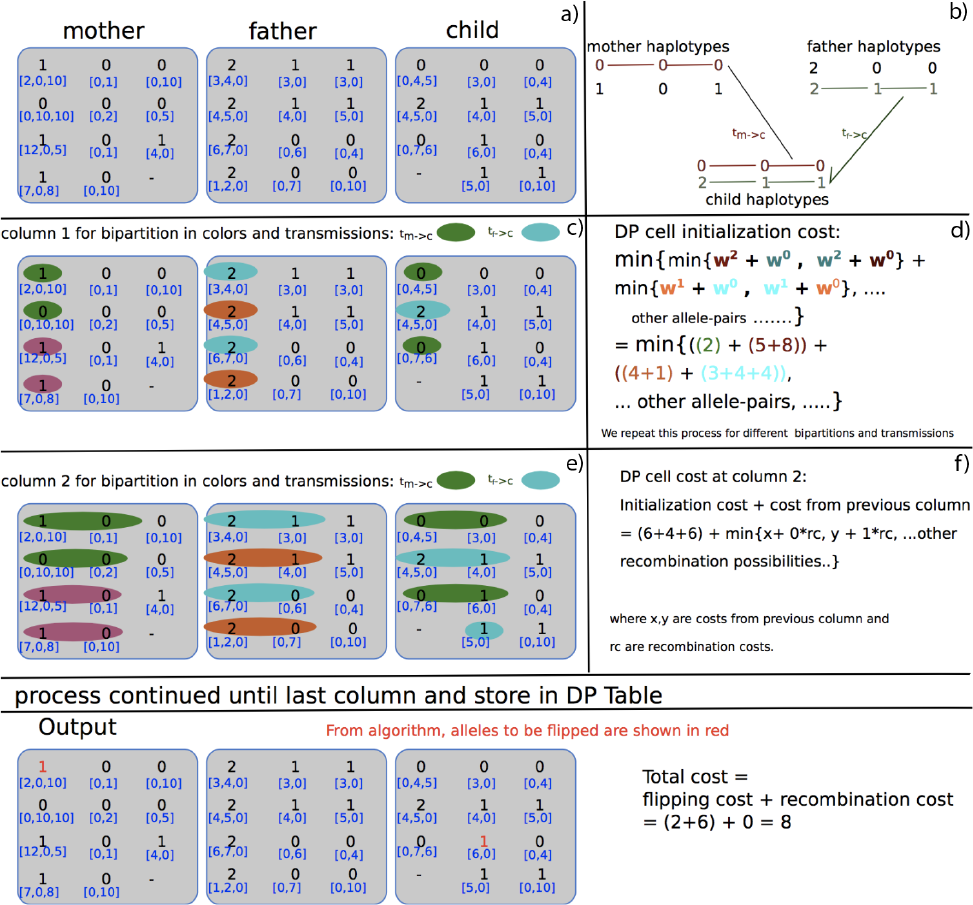
This figure shows the process of our DP algorithm. The top row shows the three input bubble matrices from mother, father and child, with their weights (a). The following rows show sample processes for DP column initialization and recurrence. In block (c), sample bipartitions and transmission values are shown with the respective calculations in block (d). The cost calculation finds the best assignment of allele-pairs to the example set of partitions which accrues the minimum cost. For example, for computing the allele assignment cost of allele 0 to the green partition, we pay a cost of 2. The other costs can be computed similarly. In block (f), the recurrence step is shown that minimizes the value of the bipartition for all previous columns, plus the initialization cost, and allowing various possibilities of recombinations. This process will be repeated for all possible transmission vectors and compatible partitions until last column. Figure adapted from Garg (2018)

#### DP column initialization

Every entry *C*(1, *B*, *t*) in the first column of the DP table is filled with the the corresponding cell initialization cost Δ_*C*_ (*k, B, t*) for all bipartitions *B* and transmissions *t*. Thus, on input the set of reads covering *l*_1_, each *C*(1, *B*, *t*) is calculated according to Equation 3.

As described in Patterson *et al.* (2014) and Garg *et al.* (2016), we enumerate over bipartitions in Gray code order, thus ensuring a runtime of 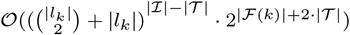.

#### DP recurrence

In the recurrence step, we compute *C*(*k, B, t*) at column *k*, which intuitively represents the optimal allele-pair assignment cost to bipartitions *B* until column *k*. In general, the cell *C*(*k, B, t*) can be computed as the optimal accumulated cost of the bipartitions *B* until *k* − 1 columns plus Δ_*C*_ (*k, B, t*) under various possibilities of recombination. Thus we add Δ_*C*_ (*k, B, t*) to values from column *k* − 1, where possible recombination costs are incurred according to the various values of *t*. The additional constraint is that only entries in column *k* −1 whose bipartitions are *compatible* with Δ_*C*_ (*k, B, t*) are considered, where two bipartitions are deemed *compatible* if they share the same readsets. By only considering compatible readsets, we are ensuring that we maintain the reality that a single PacBio read cannot come from more than one haplotype. For two bipartitions *B* and *B’*, compatibility is denoted as *B* ≃ *B’*.

In more formal terms, the value of *C*(*k, B, t*) can be written as following:

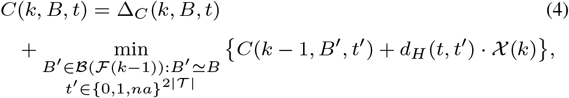

In this equation, the term *d*_*H*_(*t*, *t’*) refers to the distance, or number of changes, between two transmission vectors and thereby represents the number of recombination costs which need to be considered. In the case where any entry is *na* we assume no recombination cost.

For a sample walkthrough calculation of DP recurrence, we again turn to Figure 6. The initialization cost of the sample cell Δ_*C*_ (2, *B*, *t*) is (6+4+6) = 16, as determined according to Equation (3). In order to find the minimal value for the DP cell cost in column 2, we need to consider the costs from column 1. In the calculations shown in part (f), we only consider partitions which agree on the readsets involved, but want to consider the possibilities of recombinations between the first and second bubbles. Minimizing over all possible previous transmission vectors and compatible bipartitions gives us our DP cell value *C*(2, *B*, *t*).

We continue this process for all DP cells at column 2. In a similar manner, we repeat the DP algorithm until the last column.

#### Backtracing

At the end of our DP algorithm, the minimial value stored in the final column represents the lowest possible cost incurred. We can backtrace through the DP table to find the transmission vectors, bipartitions of readsets and final haplotype paths that led to this value.

### Generation of final assemblies

One we obtain the partitions of long-read data for each individual, we can perform haplotype-aware error-correction on the reads. Subsequently, we can use an external assembler to assemble these partitions separately create final haplotype-resolved assemblies for all individuals in a pedigree.

## 3 Datasets and Experimental Setup

Our objective was to study trios which present different genomic heterozygosity rates, and for which Illumina and PacBio data are readily available. Due to the lack of sufficient parental long-read data to pursue this goal (which major sequencing efforts will likely produce in the near future), we primarily considered simulated data for our comprehensive study of assembly behavior at varying heterozygosity rates and long-read coverage. Subsequently, we applied our method to real data of the widely-studied Col-0 and CVI-0 strains of *Arabidopsis Thaliana*.

To generate our simulated data, we created a series of diploid genomes by adding mutations at varying rates to the haploid yeast strain DBVPG6765 (Yue *et al.* (2017)). The four parental haplotypes were generated in this way; subsequently, one haplotype from each parent was selected to form the child diploid genome.

We performed this process three times to generate trio of pseudo-diploid yeast genomes with heterozygosity rates of 0.5%, 1.0%, and 1.5%. For each genome in each trio, we simulated Illumina data with average read length of 150 bp and 30× coverage. Furthermore, we simulated PacBio data for each individual in each trio at 5×, 10×, and 15× coverage. For the three trios of heterozygosity rates 0.5%, 1.0%, and 1.5%, the average PacBio read lengths, respectively, were: 6,202, 6,220 and 6,212, at 5× coverage, 6,202, 6,157 and 6,256 at 10× coverage, and 6,231, 6190 and 6,220 at 15× coverage. For comparison with TrioCanu (Koren *et al.* (2018)), we simulated child PacBio data at 15, ×, 30×, 45× and 80× coverage.

Additionally, we considered data of *Arabidopsis Thaliana*, with heterozygosity rate of 1.36%, for which assemblies have been produced by TrioCanu (Koren *et al.*, 2018). Due to unavailability of Illumina data for the Col-0 and CVI-0 strains, we simulated Illumina and PacBio data for each of them using properties of real data (Chin *et al.*, 2016). Thus, for this experiment, we were able to consider real child data and combined it with simulated parental data. We down-sampled the data for each individual in a trio to 15× coverage, with an average read-length of 17 kbp.

### 3.1 Pipeline implementation

We used a modified version of SPAdes v3.10.1 (Bankevich *et al.*, 2012) to construct our pedigree graph based on Illumina data from all related individuals. In order to maintain heterozygosity information in our graph, we ignored the bubble removal step, running it with default parameters along with the `-–only-assembler` flag, thereby producing our pedigree graph, without any error correction. Then, using VG (Garrison *et al.*, 2017), we converted the assembly graph to a *bluntified* sequence graph—that is, with redundant node sequences removed. Subsequently, we detected regions of heterozygosity, (i.e. *snarls*) with the snarl decomposition algorithm from VG (Paten *et al.*, 2017). Using GraphAligner^1^, we aligned PacBio reads from all individuals to the generated graph. Using our own implementation, we obtain bubble chains from the combined PacBio alignments according to the algorithm described in Section 2. Taking the resulting ordered bubble chains and long-read PacBio alignments, we solved the gPedMEC problem. In our calculations, we assumed constant recombination costs 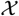 and weights in the weight matrices 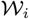 for all individuals. We determined the optimal partitions for each individual via backtracing, as detailed in our description of the algorithm. The final haplotigs were generated by assembling these computed partitions separately using Canu v1.8 Koren *et al.* (2017). These steps have been implemented as a new available tool, WHdenovo.

### 3.2 Assembly performance assessment

We evaluate the child’s predicted haplotype-resolved assemblies by aligning our predicted assemblies to the true simulated genome in our yeast-based experiments, and to TrioCanu’s published assemblies of *A. Thaliana* when handling real data. Our measures of success are introduced and described below:

#### Partitioning accuracy rate

For simulated data, we computed partitioning accuracy rate by directly comparing our predicted read partitions to the truth.

#### Average Percent Identity

We consider the best assignment of each haplotig to either of the two true references, obtained by aligning the haplotig to them. For each whole diploid assembly, we compute the average of the best-alignment percent identities over all haplotigs.

#### Assembly contiguity

We assess the contiguity of the assemblies by computing the N50 of haplotig size.

#### Assembly completeness

We assess the completeness using the total length of haplotigs assembled by each method.

## 4 Results

We present the results of our analysis of the child diploid assemblies based on the datasets described above, as assembled by both our method and by TrioCanu.

### Coverage and heterozygosity analysis

To explore a cost-effective method of assembling the child diploid genome when trio information is available, we consider PacBio datasets of varying coverage for each individual in a trio — specifically, 5×, 10× and 15× coverages. We compared our own results using coverages of 5×, 10× and 15× for all individuals in a trio to the results of TrioCanu using child PacBio data at 15×, 30×, 45×, and 80× coverages. In general, we found that TrioCanu requires 80× child coverage to achieve results comparable to our method at 15× for all individuals.

Table 1 reports the assembly performance statistics of our method applied to genomes of varying heterozygosity rates. To assess the accuracy of the child assemblies, we computed the average percent identity over both child haplotypes at varying PacBio coverages. In our approach, we observe that as we increase the long read coverage from 5× to 15× for each individual, the average identity of the haplotigs increases from 99.3% to 99.9%. This behavior is consistent across genomes of different heterozygosity rates. Table 1 also reports the performance of TrioCanu assemblies at 45×and 80× child coverage (lower coverages produced results equivalent or worse than 45×, and were hence omitted). TrioCanu produced haplotigs with average identity of 88-90% at coverages of 15×, 30×, and 45× for child data, requiring 80× to attain success comparable to our method. For real data from *Arabidopsis Thaliana* we produced haplotigs with 99.9% average identity. We believe that the Illumina-based graph used in our approach helps lead to this result. Furthermore, optimally solving the gPedMEC formulation of the phasing problem likely contributes to generating accurate haplotigs. Overall, our analysis supports the conclusion that our approach delivers accurate haplotype sequences even at a long read coverage as low as 15× for each individual in a trio.

**Table 1.** Phased assembly performance of child averaged over both haplotypes from our approach (WHdenovo) and TrioCanu. Please note that the PacBio coverage is for every individual in a trio for WHdenovo, whereas the coverage is for only child in TrioCanu.

| Approach | Het rate | PacBio cov. | #phased bubbles | %partition accuracy | Identity [%] | N50 | Total length |
| --- | --- | --- | --- | --- | --- | --- | --- |
| Child phased assemblies |  |  |  |  |  |  |  |
| Simulated data |  |  |  |  |  |  |  |
| WHdenovo | 0.5 | 3*5× | 55,561 | 94.2 | 99.3 | 21k | 7.6m |
| WHdenovo | 0.5 | 3*10× | 56,233 | 94.7 | 99.4 | 220k | 11.5m |
| WHdenovo | 0.5 | 3*15× | 56,563 | 94.9 | 99.9 | 720k | 11.9m |
| TrioCanu | 0.5 | 1*45× | — | — | 89.1 | 8.5k | 51.1m |
| TrioCanu | 0.5 | 1*80× | — | — | 99.9 | 730k | 11.9m |
| WHdenovo | 1.0 | 3*5× | 45,418 | 95.1 | 97.8 | 21k | 7.6m |
| WHdenovo | 1.0 | 3*10× | 46,228 | 96.3 | 99.4 | 210k | 11.5m |
| WHdenovo | 1.0 | 3*15× | 47,528 | 96.4 | 99.8 | 540k | 11.9m |
| TrioCanu | 1.0 | 1*45× | — | — | 88.8 | 8.6k | 51.9m |
| TrioCanu | 1.0 | 1*80× | — | — | 99.9 | 542k | 11.9m |
| WHdenovo | 1.5 | 3*5× | 35,027 | 97.4 | 97.5 | 20k | 7.5m |
| WHdenovo | 1.5 | 3*10× | 39,021 | 98.3 | 99.4 | 210k | 11.5m |
| WHdenovo | 1.5 | 3*15× | 41,126 | 98.6 | 99.8 | 520k | 11.9m |
| TrioCanu | 1.5 | 1*45× | — | — | 88.8 | 8.8k | 51.5m |
| TrioCanu | 1.5 | 1*80× | — | — | 99.9 | 540k | 11.9m |
| Real data (*A.thaliana*) |  |  |  |  |  |  |  |
| WHdenovo | 1.36 | 3*15× | 298,519 | — | 99.9 | 6.1m | 119m |
| TrioCanu | 1.36 | 1*45× | — | — | 88.9 | 81k | 400m |
| TrioCanu | 1.36 | 1*100× | — | — | 99.9 | 7m | 120m |

In measuring partitioning accuracy of long reads, we considered reads to be classified only if covering a fixed threshold of bubbles. We observe that the partitioning accuracy improves with the heterozygosity rate. For example, for genomes with heterozygosity rate of 1.5%, our calculated partitioning accuracy rate is 98.6% for 15×-fold child data. By contrast, if the heterozygosity rate is low, at 0.5%, our partitioning accuracy is 94.9% at 15×-fold data of child.

With an increase in average PacBio coverage from 5× to 15×, the haplotype contiguity achievable using our approach dramatically improves from 21 kbp to 720 kbp for trios with heterozygosity rate of 0.5%, approaching the contiguity of chromosomal-scale assemblies. When heterozygosity rate is high (>=1.0) our assemblies are somewhat fragmented (e.g. 540 kb) even at 15× coverage. This fragmentation is a result of repetitive and highly diverging regions, which cause assemblers to break contigs. For *Arabidopsis Thaliana*, we produced haplotigs with 6.1 Mb N50 length. In comparison, TrioCanu produced N50 of length 7-9 kb at coverage of 45× for different heterzygosity rates of yeast simulated data and 81 kb for *Arabidopsis Thaliana*. Using whole high-coverage PacBio data, TrioCanu was able to produce high quality assemblies; these results indicate that TrioCanu is not designed for the low coverages which our method utilizes.

Regarding haplotype completeness, our approach yields average child diploid assemblies of length ∼11.5 Mbp at 10× and 15× coverages. For real data from *Arabidopsis Thaliana*, we can produce complete assemblies of 119 Mb at 15× coverage for each individual in a trio.

In summary, our approach delivered higher quality haplotypes from 15× long-read coverage of all individuals in a trio than TrioCanu at 45× coverage of the child. The results from these experiments indicate that our approach is generalized to produce phased assemblies for genomes with different heterozygosity rates. Further, our graph-based approach is also generalized to produce assemblies for both parents (in addition to that for the child), and can help find recombination maps.

### Variant detection

As a second goal, we aimed to study haplotype-resolved variant detection. To pursue this, we aligned our predicted haplotype-resolved assemblies to each other and detected the variants, such as SNVs. From Figure 7, we observe that the the number of predicted SNVs or short indels rises in response to increasing heterozygosity rate; for example, 58,541 and 178,743 for genomes with heterozygosity rates of 0.5% and 1.5%, respectively. This result is expected because the number of variations between two haplotypes directly depends on heterozygosity rate. Additionally, the plot indicates that the number of variants we can detect with our approach (red) is very close to the true number of variants (blue). In summary, this plot indicates that our haplotype-resolved assembly approach helps to detect variants.

**Fig. 7.**
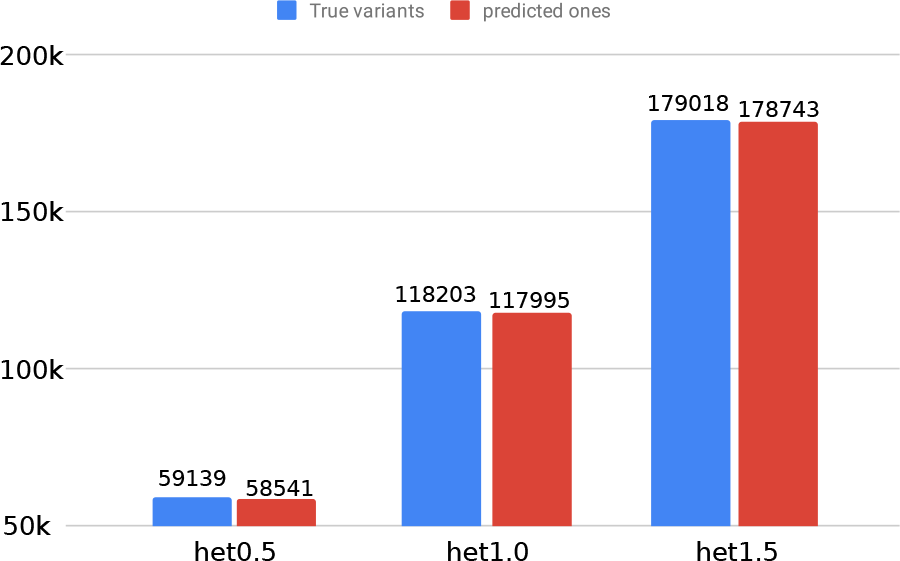
This figure shows the true and predicted variants from the phased assemblies generated using our method at various heterozygosity rates.

## 5 Discussion

Advances in sequencing technologies such as PacBio, ONT and others, which can span multiple heterozygous variants, have enabled the reconstruction of accurate phased assemblies for related individuals. Furthermore, the generation of accurate, phased assemblies for all individuals in a pedigree is essential for studies of intra-specific variation, chromosome evolution, and allele-specific expression.

The TrioCanu method (Koren *et al.*, 2018) is a hybrid approach that takes advantage of parental Illumina data and long reads from the child in a trio; yet, it has the limitation of not phasing variants that are heterozygous in all individuals. Moreover, it requires high coverage of long-read child data to produce high quality assemblies of 99.99% average identity and chromosomal-scale continuity. As sketched in Figure 2, given long reads from multiple individuals in a trio, we can efficiently phase all variants over the genome and generate chromosomal-scale assemblies.

We have developed a novel pedigree graph based approach to the problem of diploid genome assembly for pedigrees that combines short-and long-read sequencing technologies. By combining the accuracy of short reads with the contiguity offered by long reads, along with pedigree information, our approach produces accurate, complete and contiguous haplotypes. By requiring relatively low long-read coverage, our method is also a cost-effective way of generating high quality diploid assemblies. Furthermore, by performing phasing directly in the space of a pedigree graph, we can detect and phase all variants, including those that are heterozygous in all individuals. We tested our approach using simulated data from a trio of pseudo-diploid yeast genomes and real data from *Arabidopsis Thaliana*, resulting in accurate, complete haplotigs.

One restriction in our model is the use of constant recombination rates; we aim to fine tune this parameter in the future according to genomic distances, and properly incorporate recombination hotspots. Moreover, we have produced haplotype partitions of reads, but still rely on using an external assembler for producing final assemblies. The next step for this work involves producing assemblies alongside the process of graph-based phasing on pedigrees. This task may also involve phasing repeat regions, which we plan to perform using polyploid phasing as described by Chaisson *et al.* (2017). The general idea of our phasing approach can be even applied to polyploid genomes, with some gain in computation time. We will explore heuristic approaches to perform polyploidy phasing in an efficient manner, and will aim to use a joint phasing framework to obtain more contiguous diploid genome assemblies.

Our framework, in principle, is generalized to incorporate any variety of datasets; in the future, we hope to optimize our method by incorporating data such as chromatin conformation capture (Burton *et al.*, 2013) and linked read sequencing (Weisenfeld *et al.*, 2017). Each type of sequencing data will offer new information about the true haplotype paths through our graph. Additionally, incorporating PacBio CSS data (Wenger *et al.*, 2019) in our pedigree graph generation may help scale our pipeline to larger genomes. Finally, we hope to use haplotype-resolved variants we detect to shed new light on open biological questions concerning inherited disease.

## Acknowledgments

We would like to acknowledge Isaac Sebenius for writing test cases.

## Footnotes

1 https://github.com/maickrau/GraphAligner

